# A Method for Rapid Flow-cytometric Isolation of Endothelial Nuclei and RNA from Archived Frozen Brain Tissue

**DOI:** 10.1101/2021.05.12.443932

**Authors:** Amy L. Kimble, Jordan Silva, Melissa Murphy, Jessica A. Hensel, Sarah-Anne E. Nicholas, Evan R. Jellison, Bo Reese, Patrick A. Murphy

## Abstract

Endothelial cells are important contributors to brain development, physiology, and disease. Although RNA sequencing has contributed to the understanding of brain endothelial cell diversity, bulk analysis and single-cell approaches have relied on fresh tissue digestion protocols for the isolation of single endothelial cells and flow cytometry-based sorting on surface markers or transgene expression. These approaches are limited in the analysis of the endothelium in human brain tissues, where fresh samples are difficult to obtain. Here, we developed an approach to examine endothelial RNA expression by using an endothelial-specific marker to isolate nuclei from abundant archived frozen brain tissues. We show that this approach rapidly and reliably extracts endothelial nuclei from frozen mouse brain samples, and importantly, from archived frozen human brain tissues. Furthermore, isolated RNA transcript levels are closely correlated with expression in whole cells from tissue digestion protocols and are enriched in endothelial markers and depleted of markers of other brain cell types. As high-quality RNA transcripts could be obtained from as few as 100 nuclei in archived frozen human brain tissues, we predict that this approach should be useful for both bulk analysis of endothelial RNA transcripts in human brain tissues as well as single-cell analysis of endothelial sub-populations.

## Introduction

Endothelial cells line the vasculature of the brain, where they contribute to the function of a wide range of fundamentally important tissues including maintenance of the blood brain barrier, immune cell surveillance, blood flow, and the maintenance of neural stem cells. Despite their critical brain functions, endothelial cells are a relatively small proportion of all cells in the brain, with ∼2-4% in total area coverage ^1^, and ∼0.2-7% as a total proportion of all brain cells based on single-cell sorting experiments ^2–6^. As a result, single-cell experiments using whole brain tissue have lacked sufficient resolution of the endothelial cell population for reliable transcriptional analysis ^3^. More focused approaches, based on the specific isolation of endothelial cells have provided better transcriptional analysis in bulk ^7,8^ and in sorted single cells ^9^. However, these tissue digestion protocols are complicated by a balance between the time in harsh 37°C digestion conditions needed to release single cells and the damaging effects of these conditions on RNA profiles and cell-surface markers. Digestion-insensitive fluorescent reporters have been useful ^9^, but only in mouse models. Together with the limited availability of acutely isolated human brain samples, these complications have severely limited the analysis of endothelial cell transcripts in human brain pathology.

Nuclei isolation has emerged as a valuable addition to the toolkit for investigators interested in transcriptional responses in cells. Although there are limitations, in that the nuclear RNA pool is enriched for nascent or nuclear-localized transcripts, there is a good correlation between the nuclear RNA pool and the whole-cell RNA pool ^2,6,10,11^. Furthermore, nuclear isolation provides a number of distinct advantages. First, the isolation process is rapid, and does not require enzymatic dissociation. As the mechanical and biochemical processes needed to dissociate cells (∼ 1 hr at 37°C) can activate RNA responses, the rapid protocols involved in nuclei isolation (<5 min at 4°C) better preserve acute tissue responses ^12^. Second, although there are limitations in looking only at nuclear RNA, this can also be an advantage, as it limits the number of ribosomal and mitochondrial RNA transcripts detected, and increases the detection of new transcripts. Finally, and perhaps most importantly, nuclear isolation can be applied to frozen archived tissues. This is particularly important in human studies, where such archived tissues are typically the only samples available. Single nuclei analysis of whole brain tissues ^3^ or from microvascular fragments provides a valuable assessement of transcript levels among individual nuclei (BioRxiv papers, Heimann & Wyss-Coray labs). However, these single cell nuclei approaches are costly when applied across hundreds of samples and generally confined to the 3’ tagging approach in current single cell work flows, and thus not amenable to the analysis of full transcripts and RNA splicing.

Here, we use the expression of an endothelial cell-specific transcription factor to enrich for endothelial nuclei in archived frozen mouse brains. We show this approach enriches endothelial-specific transcripts, while depleting transcripts of non-endothelial cells, and allows analysis of RNA splicing across transcripts. We then apply this technique to a set of archived human brain tissues from the NIH NeuroBioBank, and show a similar enrichment of endothelial transcripts. We suggest that this will be a valuable method for the analysis of endothelial cell types and differential responses in model systems, like the mouse, and in human disease.

## Materials and Methods

### Tissue sources

Human embryonic kidney (HEK293) and human umbilical vein endothelial (HUVEC) cells were obtained from ATCC, and cultured in 10% FBS DMEM (HEK293) or endothelial growth kit + VEGF (ATCC). C57Bl6/J mice were obtained from JAX, and used for the isolation of brain hemispheres. Frozen human brain tissue (Brodmann area 10) was obtained from the NIH NeuroBioBank.

### Nuclei isolation and flow-cytometry

Cells (1M) and tissues (200mg) were homogenized in Nuclei EZ lysis buffer (Sigma Nuc 101) with RNase inhibitor (0.04U/uL final, Clontech Cat #2313A). Cultured cells were homogenized by trituration by pipet. Frozen mouse and human brain tissues were mechanically homogenized (Next Advance Bullet Blender BB724M with 3.2 mm stainless steel beads using setting 4 for 4min at 4C). After homogenization, nuclei were diluted to 5mL and spun down at 700xG for 5min, and then washed once 5mL Nuclei EZ lysis buffer, incubated 2min on ice before spinning down a second time at 700xG for 5min. Nuclei were then fixed in 600uL PBS + 0.5% PFA for 1min at 4C, before adding PBS with RNAase inhibitor (0.04U/uL final) and spinning down at 700xG for 3min. Cells were resuspended in 1mL of PBS with RNAase inhibitor (0.04U/uL) and BSA (0.1% w/v). Note, increased salt (e.g. 500mM NaCl) can be used to further limit RNAse activity during staining, but must be tested with antibodies to be used. Nuclei were passed over a 70um and then 35um filter. A small aliquot was taken for unstained nuclei and single color controls, and the rest spun down again at 700xG for 5min at 4C for staining with the antibody mixture in 100uL PBS + RNAse inhibitor + BSA for 15min at 4C (1:200 [2.5ug/mL final] Anti-Erg 647, Clone EPR3864, Abcam and 1:200 Anti-NeuN Cy3 Clone A60, Sigma), and DAPI for nuclei labeling (1:2000 of a 5mg/mL stock). After incubation, samples were washed with 1mL PBS + RNAse inhibitor + BSA and spun down at 700xG for 5min before resuspending in PBS + RNAse inhibitor + BSA with 0.6U/uL DNAse (to break clustered nuclei) and filtereing over a 35um filter. Nuclei were then sorted on an Aria 2 with 40um nozzle and collected into a dry tube at 4C.

### RNA-isolation and sequencing

RNA was isolated from fixed nuclei using an FFPE kit (RNAeasy, Qiagen 73504) with on column DNAse treatment. For quantitative PCR analysis, cDNA was prepared using random priming (Applied Biosystems kit). For RNA-sequencing, samples were prepared for library prepraration using the SMARTer® Stranded Total RNA-Seq Kit v2 - Pico Input Mammalian Kit for Sequencing (Takara Bio USA, Mountain View, CA). Total RNA was quantified and purity ratios determined for each sample using the NanoDrop 2000 spectrophotometer (Thermo Fisher Scientific, Waltham, MA, USA). To further assess RNA quality, total RNA was analyzed on the Agilent TapeStation 4200 (Agilent Technologies, Santa Clara, CA, USA) using the RNA High Sensitivity assay. Amplified libraries were validated for length and adapter dimer removal using the Agilent TapeStation 4200 D1000 High Sensitivity assay (Agilent Technologies, Santa Clara, CA, USA) then quantified and normalized using the dsDNA High Sensitivity Assay for Qubit 3.0 (Life Technologies, Carlsbad, CA, USA).

Sample libraries were prepared for Illumina sequencing by denaturing and diluting the libraries per manufacturer’s protocol (Illumina, San Diego, CA, USA). All samples were pooled into one sequencing pool, equally normalized, and run as one sample pool across the Illumina NextSeq 550 using version 2.5 chemistry. Target read depth was achieved per sample with paired end 150bp reads.

### Read-mapping and transcript analysis

Transripts were mapped to the mouse (mm9) or human (Hg38) genome by STAR ^13^ (2.5.3a) with parameters --outFilterType BySJout --outFilterMultimapNmax 20 --alignSJoverhangMin 8 --alignSJDBoverhangMin 1 --outFilterMismatchNmax 999 --alignIntronMin 10 --alignIntronMax 1000000 --alignMatesGapMax 1000000. Reads were filted so that only intron spanning reads were used, to remove background from DNA contamination, using samtools (samtools view -h *bam_filename*| awk ‘$6 ∼ /N/ || $1 ∼ /^@/’ | samtools view -bS - >

*spliced.bam_filename*. Gene level counts were obtained from the bam files using Whippet ^14^.

### Analysis of other data sets

Top 1000 enriched mouse and human genes were taken from Supplemental File 1 of McKenzie et al ^15^. Gene level transcript frequency estimates (FPKM) for whole brain and sorted cells from mouse and human brain were taken from the Supplemental Table 4 of Zhang et al ^16^. DropNucSeq data from Habib et al. was obtained from the Broad Institute’s Single Cell Portal developed as a part of the BRAIN (Brain Research through Advancing Innovative Neurotechnologies) initiative ^2^. Average read count (gene level) were calculated in R by cluster and for all nuclei (all brain nuclei). The endothelial cluster was confirmed by expression of key endothelial specific gene transcripts (Erg, Cldn5, Cdh5, Pecam, Icam2).

### Analysis of enrichments

Enrichment of transcripts obtained in whole sorted endothelial cells was obtained by Log2[Endothelial FPKM (fragments per million) / Whole brain FPKM], using expression values from Supplemental Table 4 of Zhang et al. Similarly, enrichment of transcripts obtained in the endothelial cluster of nuclei from mouse and human brain was obtained by Log2[Endothelial Average RPM (reads per million) / Average all brain nuclei RPM]. Finally, enrichment of transcripts in sorted Erg+ nuclei was obtained by Log2[Erg+ TPM / Whole brain FPKM] and Log2[Erg+ TPM / Whole brain nuclei avg. read count], using data from Zhang et al. (whole cell) and Habib et al. (nuclei).

## Results

### Enrichment of endothelial cells by Erg antibody staining

As prior work has used a neuronal-specific transcription factor (NeuN) to enrich neuronal nuclei from brain tissues ^17,18^, we hypothesized that we could use an endothelial-specific transcription factor to enrich endothelial nuclei from tissues. Erg is a highly specific endothelial transcription factor in mouse and human tissues (Figure 1A&B). To determine whether Erg can be used to enrich endothelial cells by flow cytometry, we isolated nuclei from human umbilical endothelial cells (HUVEC), human embryonic kidney cells (HEK293), and a mixture of the two populations. Analysis of the stained mixtures by flow cytometry showed staining of all HUVEC cells and none of the HEK293 cells in isolation (Figure 1C&D). The mixture of the cell types showed two distinct populations (Figure 1E), suggesting that endothelial nuclei could be sorted from a mixture based on expression of Erg.

**Figure 1.**
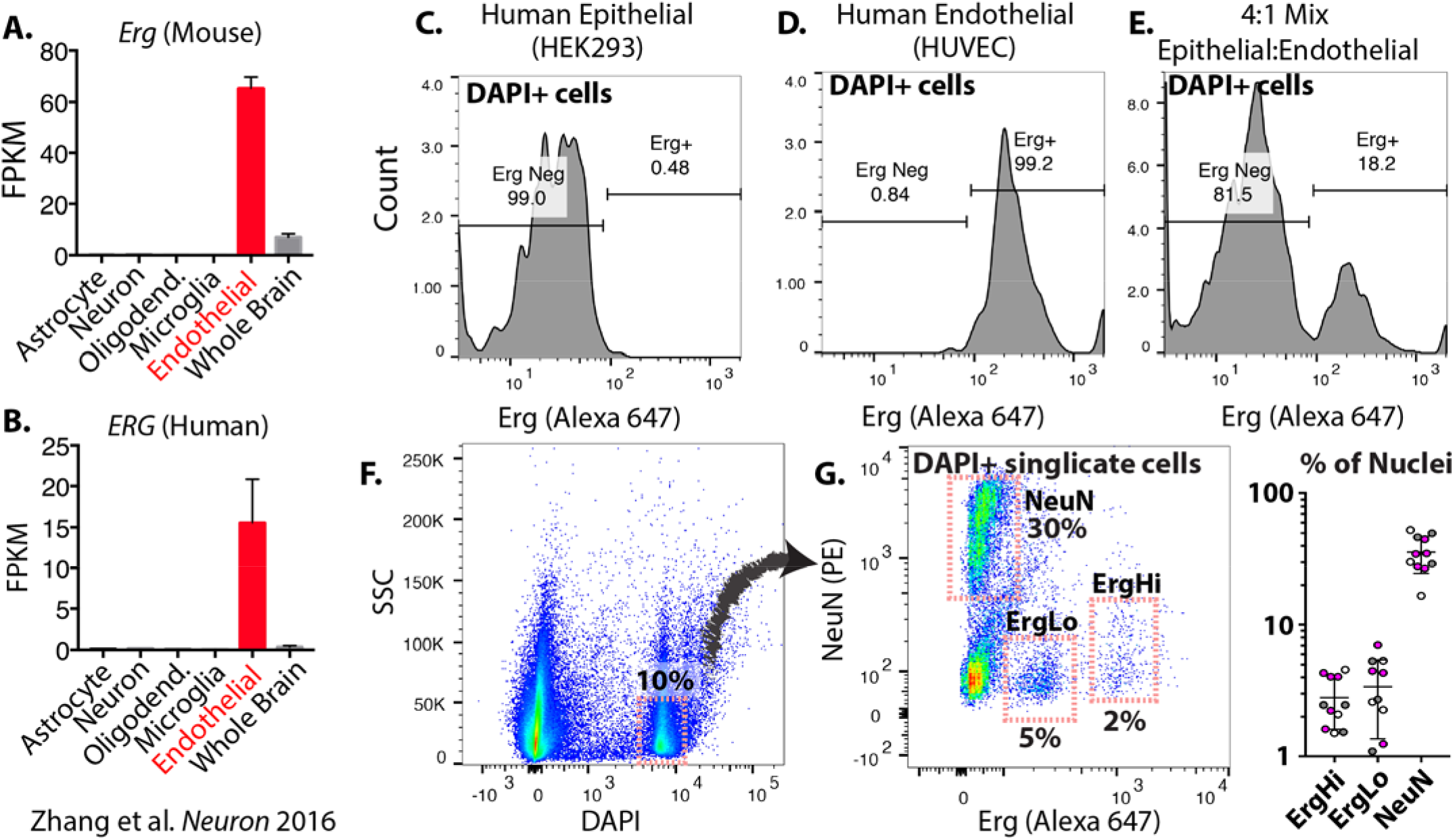
Erg marks isolated endothelial nuclei for flow cytometry. (A&B) Erg expression in the indicated cells isolated from mouse (A) and human (B) brain tissues, as previously reported (mean +/- SD) ^16^. (C-E) Flow-cytometry plots showing Erg-conjugated Alexa647 signal intensity in DAPI+ nuclei isolated from HEK293 (C), HUVEC (D), or a mixture of HEK293 and HUVEC cells (E). (F&G) Flow-cytometry plots showing DAPI+ nuclei in human brain tissue (F) and the fluorescence intensity from neuronal marker NeuN (PE conjugated) or Erg (Alexa 647 conjugated). Gates show selected neuronal and endothelial populations and the proportion of total DAPI+ nuclei. FPKM = fragments per kilobase of exon model per million reads mapped. Graph shows the percent of nuclei found in each fraction from multiple frozen human cortex samples. White dots = young unaffected brain, grey dots = old unaffected brain, purple dots = dementia brain.

To determine whether Erg staining could be applied to archived human brain tissues, we performed nuclear isolation and staining of 200 mg of brain tissues obtained from the NIH NeuroBioBank. To assess staining across a range of sample, we included tissues from young and old donors, as well as donors with documented dementia. To fully and rapidly dissociate the brain tissue and to extract nuclei, we used a mechanical homogenizer, which was combined with a nuclei isolation buffer. We found that Erg stained two subpopulations of nuclei from the human brain (ErgLo and ErgHi, Figure 1F), which were clearly distinct from the NeuN neuronal population. The number of Erg-stained nuclei (∼2%) correlates well with the expected proportion of nuclei in whole brain, based on prior whole brain single-cell and nuclei sequencing experiments [2-7]. Importantly, the method yielded reproducible results, leading to the reliable detection of 1-4% ErgHi+ nuclei and 1-8% ErgLo+ nuclei across multiple samples from the NIH NeuroBioBank (Figure 1G, and SI Table 1). Notably, this is also consistent with the amount of endothelial cells in histological sections ^1^.

These data suggest that Erg specifically stains endothelial nuclei and could allow for their isolation from frozen human brain tissues.

### Extraction of RNA from Erg+ nuclei confirms enrichment of endothelial transcripts

Our goal is to assess RNA transcripts in isolated endothelial nuclei. To benchmark this approach, we extracted nuclei from freshly frozen mouse brain tissues, and again stained for Erg and NeuN. The mouse brain tissues showed similar staining patterns and similar proportions of Erg+ and NeuN+ nuclei to the human brain tissue (Figure 2A&B). Notably, the murine brain showed only a single Erg population, not a high and low population as in the human brain (Figure 1A-C).

**Figure 2.**
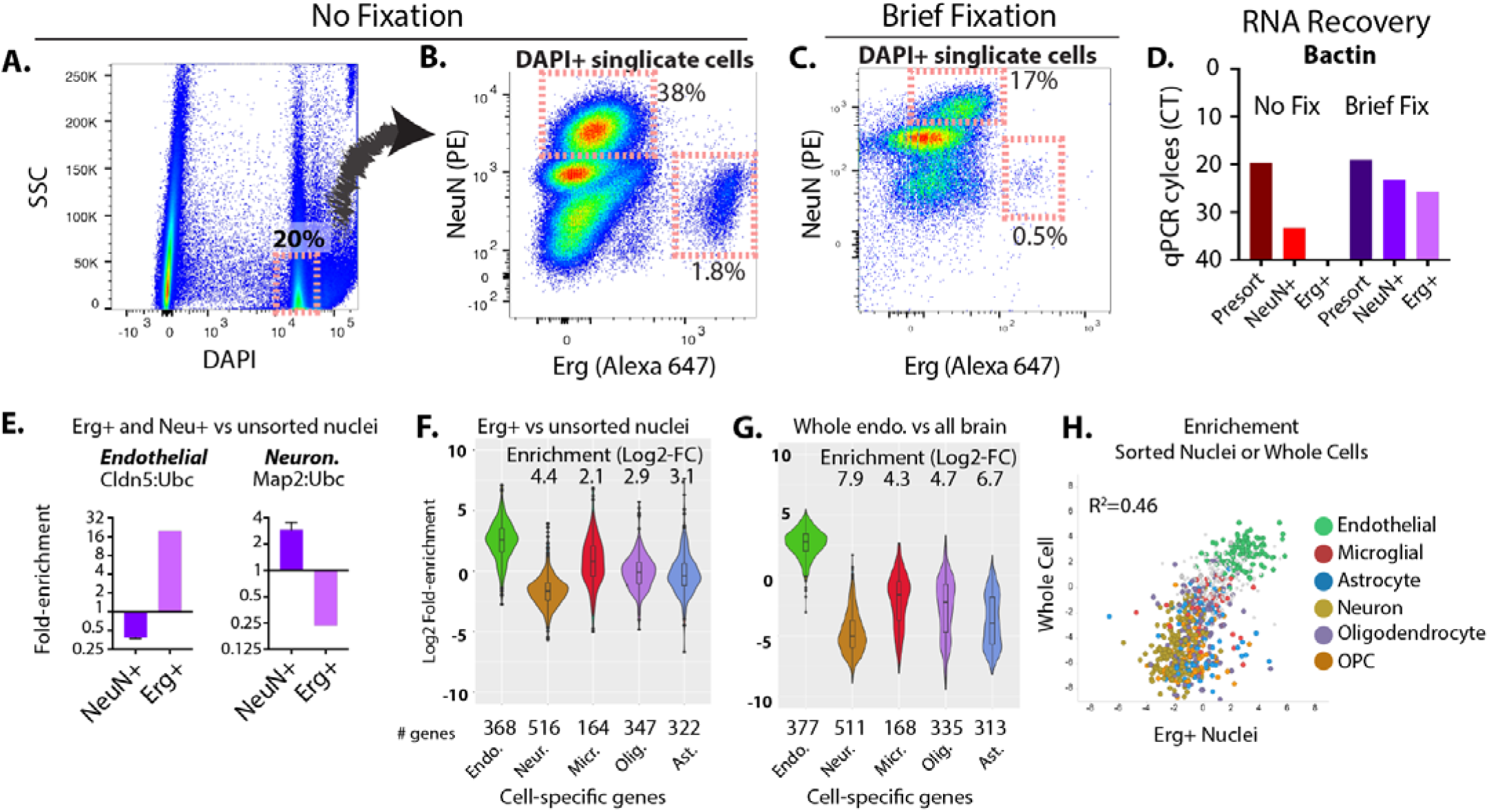
Isolation of Erg+ nuclei enriches for endothelial transcripts in mouse brain tissue. (A-C) Flow-cytometry plots showing DAPI+ and low side scatter (SSC) low nuclei in mouse brain tissue (A) and the fluorescence intensity from neuronal marker NeuN (PE conjugated) or Erg (Alexa 647 conjugated) in unfixed (B) or fixed (C) samples. Gates in (B&C) show selected neuronal and endothelial populations and the proportion of total DAPI+ and SSC low nuclei. (D) Shows the qPCR cycle threshold (CT) from unsorted starting material and the sorted equivalent populations of unfixed and fixed nuclei. A larger CT value indicates reduced amounts of RNA detected (on a Log2 scale). (E) Enrichment of cell type-specific RNA by quantitative PCR from the sorted Erg+ or NeuN+ fixed nuclei versus unsorted fixed nuclei. The fold-change increase in each transcript, relative to unsorted nuclei is shown. (F&G) Enrichment of the cell-specific RNA in endothelial cell (Erg+ nuclei, or historical data from whole sorted endothelial cells) versus all brain nuclei from RNA-sequencing analysis. Enrichment analysis shows the change in spliced RNA reads for each gene of the marker gene set, relative to unsorted nuclei. Median enrichment relative to other sets of markers is shown. (H) Plot showing the correlation in enrichment in transcripts from each gene from Erg+ nuclei versus unsorted nuclei, or whole endothelial cells (Zhang et al.) versus whole brain. Characteristic marker genes are indicated. (Endo=endothelia, Neur.=neuron, Micr.=microglia, Olig.=oligodendrocyte, Opc.=oligodendrocyte precursor cell, Ast.=astrocyte).

An obstacle that appeared was that although cells were enriched for endothelial RNA, that the relative RNA levels were low. While this was only a minor effect in cells after staining and prior to sorting, very little RNA was obtained after sorting, suggesting losses to RNA leak during the sorting process (Figure 2D and Supplementary Figure 1, compare Erg+ and NeuN+ sorted populations to unsorted population without fixation). Therefore, we performed a mild paraformaldehyde (PFA)-mediated cross-linking of RNA immediately after nuclei isolation and prior to staining and sorting, with the goal of limiting RNA leak from the cells during the sorting procedure. We found that Erg+ nuclei could be isolated after this brief cross-linking (Figure 2C), although the proportions of each labeled population was reduced. Importantly however, fixation yielded much high levels of RNA in the sorted nuclei (Figure 2D and Supplementary Figure 1, compare Erg+ and NeuN+ sorted nuclei with and without fixation).

To determine whether our fixation protocol and Erg+ and NeuN+ sorting enriched for endothelial- and neuronal-specific transcripts, we examined expression of endothelial- and neuronal-specific markers in RNA from both the unsorted nuclei and the sorted nuclei by quantitative PCR (qPCR). We found a ∼20-fold enrichment in the endothelial marker Cldn5 in the Erg+ nuclei (relative to Ubc, a housekeeping gene), and a depletion of these markers in the NeuN+ nuclei (Figure 2E). In contrast, we saw a depletion in neuronal marker Map2 in Erg+ nuclei and a ∼3-fold enrichment in this marker in NeuN+ sorted nuclei (Figure 2E). As endothelial cells are ∼1/20 of the total brain nuclei and neuronal cells are ∼1/3 of the total brain nuclei, these relative enrichments suggested nearly optimal enrichment of RNA using these nuclear markers.

To broadly examine enrichment of endothelial- and neuronal-specific transcripts, we performed RNA sequencing on the isolated RNA from Erg+ and NeuN+ nuclei. From material derived from ∼2.5K sorted Erg+ nuclei, RNA was made into a cDNA library using the SMARTer v2 stranded prep kit. As previously reported [2, 7, 11, 12], we found a high number of intronic reads. We also found a large fraction of intergenic reads, likely due to DNA reads, and possibly a consequence incomplete DNAse digestion after fixation . To focus on only RNA transcript, we confined our analysis to splice junction spanning reads and detected transcripts from 15.5K unique genes.

To determine whether endothelial transcripts were in fact enriched, and to see whether other cell populations might also be enriched (due to promiscuous binding of the Erg marker, for example), we looked for the enrichment of RNA markers of specific cell types based on a meta-analysis of single-cell data from mouse brain samples [8]. We used the top 1000 genes enriched in each cell type: endothelial, microglial, astrocyte, neuronal, oligodendrocyte, and oligodendrocyte progenitor cell (OPC). To control for possible differences in the enrichment of these markers in nuclei versus whole cells, we plotted these markers on two gold-standard datasets, the whole cell analysis conducted by Zhang et al. [8], and the nuclei analysis (DropNucSeq) conducted by Habib et al. [2], examining enrichment of endothelial markers and depletion of other markers in a comparison between endothelial cell transcripts versus bulk brain (SI Figure 2A). Using this panel of markers, with minor modifications to improve the stringency of endothelial markers (SI Figure 2C), we looked for enrichment of gene transcripts in Erg+ sorted nuclei versus unsorted brain nuclei. We found that these endothelial-specific genes were enriched on average by ∼3 (Log2) versus other non-endothelial markers, or ∼8-fold (Figure 2F). Notably, the best enrichment was against neuronal nuclei (∼4.4 (Log2), or 20-fold), which were negatively depleted from the Erg+ population by the NeuN marker. Enrichment was comparable, but diminished, relative to whole endothelial cell nuclei (Figure 2G). No other cell-specific markers were similarly enriched, supporting the specificity of endothelial cell nuclei isolation by Erg+ staining. Plotting the relative enrichments in transcripts in Erg+ sorted nuclei (relative to unsorted nuclei) against those previously published data from whole endothelial cells (relative to bulk brain tissue), the group of endothelial specific transcripts is similarly enriched in both (Figure 2H).

Our data suggest that Erg+ allows the efficient isolation of endothelial transcripts from whole frozen mouse brains.

### Extraction of RNA from Erg+ nuclei in frozen archived human brain tissue

Our ultimate goal has been to develop a method able to assess RNA transcripts in human brain endothelial cells in frozen archived samples. Therefore, we used the protocol we had benchmarked on mouse brain tissue to examine enrichment of endothelial RNA in Erg+ human brain nuclei, using a panel of human cortical samples from the NIH NeuroBioBank (SI Table 1). We isolated RNA from fixed and sorted nuclei, and performed quantitative PCR using endothelial- and neuronal-specific markers. Unlike the mouse brain, there were two Erg+ populations in the human brain, high and low; we examined each separately. We found that, relative to housekeeping transcript UBC, endothelial transcript CDH5 was enriched ∼30-fold from unsorted nuclei in the ErgHi+ nuclei, but not NeuN+ nuclei (Figure 3A). In contrast, the neuronal marker MAP2 was enriched ∼5 fold in the NeuN+ nuclei, but not the Erg+ nuclei (Figure 3A). To confirm that isolated nuclei were only from the vascular endothelium and not mural cells adjacent to the endothelium (pericytes or smooth muscle cells), we also examined enrichment of transcripts specific to those cell types. We found that neither PDGFRB (pericyte marker) nor ACTA2 were enriched in the Erg+ or NeuN+ nuclei, relative to the unsorted nuclei (SI Figure 3). Thus, enrichment of Erg+ nuclei in the human brain appeared to have performed as well or better than enrichment from the mouse brain.

**Figure 3.**
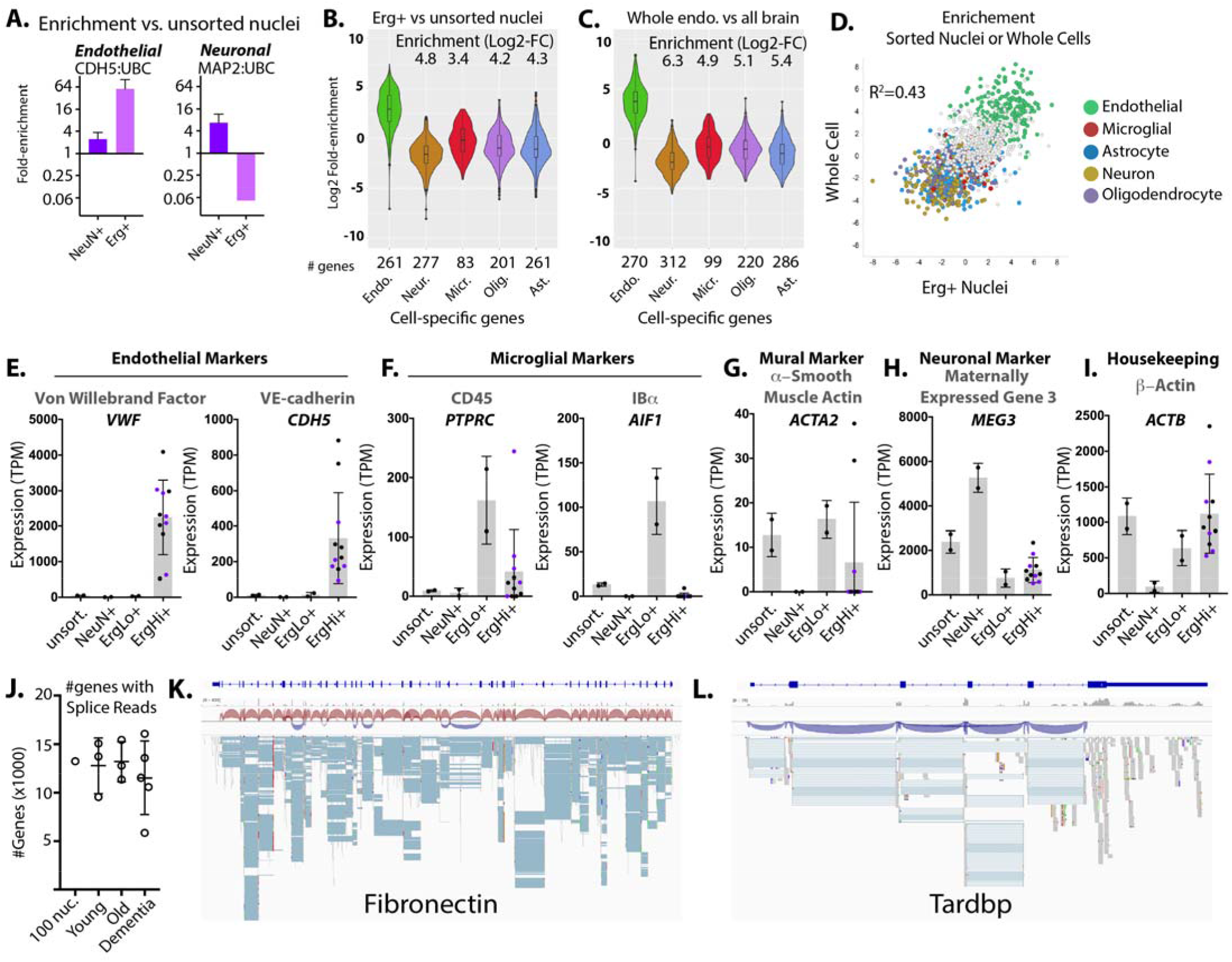
Isolation of Erg+ nuclei enriches high-quality endothelial-specific transcripts from archived human brain tissues. (A) Enrichment of cell type-specific RNA from the sorted Erg+ or NeuN+ nuclei versus unsorted nuclei. The fold-change increase in each transcript, relative to unsorted nuclei is shown. (B&C) Enrichment of the cell-specific RNA in endothelial cell (Erg+ nuclei, or published data from whole sorted endothelial cells ^16^) versus all brain nuclei from RNA-sequencing analysis. Enrichment analysis shows the change in spliced RNA reads for each gene of the marker gene set, relative to unsorted nuclei. Median enrichment relative to other sets of markers is shown. (D) Plot showing the correlation in enrichment in transcripts from each gene from Erg+ nuclei versus unsorted nuclei, or whole endothelial cells versus whole brain ^16^. Characteristic marker genes are indicated. (E-I) Gene expression (transcripts per million, TPM) of reads spanning splice junctions for the indicated genes showing (E) endothelial markers, (F) microglial markers, (G) mural cell markers, (H) neuronal markers, and (I) a housekeeping gene. Samples from dementia donors are indicated in purple. (J) Number of genes with reads spanning splice junctions. (K&L) Examples of reads plotted against the genome, from STAR alignment. The organization of introns and exons is shown in dark blue above. Plot shows splice junction spanning reads (blue) and reads (grey). Direction of the reads is shown in the upper panel sashimi plot (red, from right to left and blue, from left to right). (Endo=endothelia, Neur.=neuron, Micr.=microglia, Olig.=oligodendrocyte, Opc.=oligodendrocyte precursor cell, Ast.=astrocyte).

To examine this more broadly, we generated cDNA libraries and performed RNA sequencing on sorted ErgHi+, ErgLo+, NeuN+ and unsorted nuclei from the panel of samples using the SMARTer v2 stranded prep kit, and performed 100M 150bp reads on each sample. We examined the top cell-specific transcripts enriched in major brain cell types, as we had done in the mouse brain, by focusing on only splice junction reads (Figure 3B-D and SI Figure 2). Again, we found enrichment of endothelial cell transcripts in the Erg+ population (Figure 3B) which closely resembled the enrichment of these same transcripts in whole endothelial cells (Figure 3C). We saw better enrichment in human tissues than in mouse brain (4.8 vs 4.4 Log2 enrichment vs neuronal transcripts, and 3.4 vs 2.1 Log2 enrichment vs microglial transcripts; see Figure 2F and 3B). As in mouse brain, microglial markers were least depleted, but one possibility is that this is due to substantial overlap in endothelial and microglial genes. Transcripts highly enriched in whole endothelial cells were generally also enriched in ErgHi+ nuclei (Figure 3D).

To examine enrichment among the different samples in the panel, we calculated transcript levels, based on transcripts per million (TPM) using splice junction reads (Figure 3E-I). We found that characteristic endothelial transcripts Von Willebrand Factor (VWF) and VE-cadherin (CDH5) were highly enriched in the ErgHi+ nuclei (Figure 3E). In contrast, microglial transcripts CD45 (PTPRC) and IBa (AIF1) were enriched in the ErgLo+ nuclei (Figure 3F), suggesting that microglia express lower, but detectable levels of Erg. Notably, the mural cell marker smooth muscle α−actin (ACTA2) is not enriched in the Erg+ fraction (Figure 3G), and neuronal marker MEG3 is enriched in neuronal cells but not ErgHi+ nuclei (Figure 3H). Housekeeping gene beta-actin (ACTB) is expressed similarly in all population except neuronal cells, where actin expression is predicted to be low (Figure 3I). Thus, an analysis of the most tissue-specific canonical markers supports the high enrichment observed in bulk marker analysis.

We obtained an average of 12.3K +/- STDEV 2.3K transcripts in each Erg+ population analyzed (Figure 3J). This allowed a level of splice isoform analysis not possible from poly-A primed annotation (e.g. droplet methods of single cell analysis). To test the limits of this, we examined read coverage of two large and alternatively spliced transcripts important in endothelial cell biology in a sample obtained from 100 sorted nuclei (Figure 3K&L). We obtained coverage of both constitutive and alternatively spliced junctions in both Fibronectin (Figure 3K) and Tardbp (Figure 3L), showing that splicing analysis is possible from as few as 100 sorted Erg+ nuclei.

Thus, our data suggests that Erg efficiently isolated endothelial cell nuclei from archived frozen human brain, as it did from snap-frozen mouse brain tissues, allowing for the detailed analysis of both total transcript levels and isoforms.

## Discussion

Here we establish a novel approach for the enrichment and analysis of endothelial RNA transcripts from frozen archived brain tissues based on nuclei sorting by the endothelial transcription factor Erg. Our approach selectively enriches endothelial transcripts from whole brain nuclear homogenate, reproducibly enriching nuclei with expression of endothelial transcripts (e.g. VWF and CDH5) several hundred times over unsorted nuclei among a diverse set of frozen cortical tissues. We show that this approach can be coupled with RNA sequencing using as few as 100 sorted nuclei from archived human brain tissues. As blood brain barrier dysfunction is a major contributor to neurodegenerative human diseases, the ability to specifically interrogate the endothelial cells in archived human tissue samples will provide a new perspective on these diseases.

### Erg as a marker of endothelial nuclei

The specificity and sensitivity of Erg as a nuclear marker of endothelial cells is critical in our approach. Analysis of both mouse and human brain tissues has shown that high levels of Erg are nearly exclusively expressed in endothelial cells (Figure 2&3). Antibody staining of both mouse and human tissues has confirmed the specificity of this transcription factor to endothelial cells within brain tissue, and shown that nearly all endothelial cells are positively stained ^19–21^. While endothelial-to-mesenchymal-like transitions in endothelial cells in tumors have been linked to reduced Erg expression ^22^, we did not observe a substantial reduction in Erg expression, or in the isolation of endothelial transcripts in the ErgHi+ population in any of the human samples we have examined. While this does not exclude the possibility that a small fraction of endothelial cells may become ErgLo or negative under some conditions, our own in vivo data does not suggest a loss of Erg transcript following activation in vivo or in vitro ^23^. Thus, in our hands, and across a range of human cortical specimens, high levels of Erg expression is a reliable indicator of endothelial cell nuclei. This should be differentiated from nuclei with a low level of Erg expression (ErgLo), which we find exhibit markers of microglia and not endothelial cells (Figure 3).

Notably, Erg appears to be a good endothelial cell marker in a variety of other tissues outside of the brain, suggesting that the general approach that we have outlined here may also be useful in rapidly isolating endothelial nuclei from these tissues as well.

### Advantages and limitations of nuclei-based RNA isolation

The primary advantage of nuclei isolation is that RNA can be isolated from individual cells of frozen tissues, as the nuclei remain relatively intact after freeze-thaw, while the cytoplasmic membranes of large cells, like endothelial cells and neuronal cells, are severely disrupted and typically not recoverable. We again confirm that nuclei isolation is a very efficient process for the isolation of cell-specific RNA from frozen tissues, using both rapidly snap-frozen mouse brain tissue and a typical archived human brain sample. Our nuclei isolation approach is extremely rapid (<5 min) and never exposes cells or nuclei to the 37°C incubations typically used for the isolation of whole cells. As prolonged cell digestion procedures have been shown to obscure acute response genes in neuronal cells, we applied this rapid nuclei isolation protocol to mouse models, as well as the archived human brain tissues that were our first target.

Despite the advantages of rapid isolation of nuclei, there are also disadvantages that should be noted. First, the nucleus contains only some of the cells’ RNA. However, this is still a substantial fraction, ∼20-50% by some estimates ^11^. The specific transcripts found in the nucleus and the cytoplasm vary, as more ribosomal RNA and fully processed mRNA encoding proteins are found in the cytoplasm while actively transcribed and intron-detained transcripts are enriched in the nucleus ^11,24^. While this might at first appear to be a disadvantage, assessment of actively transcribed and nuclear RNA is emerging as a powerful means of examining changes in cell state ^25^. Intronic reads are a critical component of this analysis, as they are used to define the levels of newly transcribed and nuclear RNA.

Second, nuclei isolation is limited by the availability of nuclear markers of specific cell types. Decades of work have established abundant cell membrane markers (CD45, CD3, CD4, CD8, CD11b, CD31, etc.) which allow for positive and negative selection of cell populations. Only a few nuclear markers have been established, NeuN (Rbfox3, a neuronal marker) and Erg, which we establish here as a reliable marker of endothelial cell nuclei by flow. However, the principles applied to NeuN and Erg could easily be applied to other cell-type specific nuclear proteins, such as Pu.1 for macrophages and microglia, and TCF-1 for lymphocytes. Combinatorial approaches, as we show here for Erg and NeuN could be used for other cell-type specific transcription factors to provide a similar positive and negative gating approach.

A final consideration in the comparison of nuclei vs whole cell isolation is the retention of RNA through the process. In our analysis of samples from freshly frozen mouse brain tissues, we found that some RNA was maintained within the nucleus through the sorting of nuclei. However, we found that cross-linking the RNA into place resulted in a substantial increase in the RNA obtained from a similar amount of input. Most of these losses occurred in the sorting of the nuclei. As large antibodies are able to enter the nucleus, this suggests that holes in the nuclear membrane produced by the nuclear extraction procedure could allow RNA leak, which may be exacerbated by large dilution and shear forces in flow cytometry. While cross-linking RNA into place provided a much better yield of RNA, it can also cause problems with longer RNA reads by leading to RNA adducts, only some of which are successfully removed ^26^. We experimented with other approaches of reversible cross-linking, including DSP ^27^, but found that very brief paraformaldehyde fixation yielded the only reliable results.

### Future applications

We were able to obtain highly enriched endothelial transcripts from only ∼100 nuclei from archived frozen human brain tissue. As we routinely obtain 5-10K Erg+ nuclei from these same archived brain tissues, and single nuclei have been used effectively for the analysis of transcripts at the single-cell level, we propose that our approach might be coupled with these single nuclei tools to enable a deep interrogation of endothelial transcriptional responses in disease states. Work on the endothelial cells of the brain using enrichment approaches at the whole-cell level have revealed distinct cell types within the endothelium, corresponding to arterial cells, venous cells, and subtypes of capillary endothelial cells ^9^. Assessment of changes within these subsets of the endothelium will be important in understanding changes in dementia and other diseases of the brain.

Another future application will be the correlations between gene expression profiles, and perhaps also chromatin accessibility by Assay for Transposase-Accessible Chromatin using sequencing (ATAC-seq), and cell state as defined by levels or phosphorylation states of nuclear transcription factors and splice factors. For example, levels of TDP-43 (Tardbp), a splice factor excluded from the nucleus in the progression of a wide range of neurodegenerative disease, has been used to sort neuronal cell populations ^28^. We have shown that this marker can also be assessed in endothelial cells (Figure 3F). By adding barcoding oligomers to antibodies, approaches like CITE-seq allow for the assessment of hundreds of proteins or protein modifications in single cells ^29^. These approaches may be combined to correlate the levels of nuclear factors with RNA transcripts and splicing activity.

Using a rapid nuclei isolation protocol and Erg as an endothelial cell marker, we were able to effectively sort brain endothelial cell nuclei from frozen mouse and archived human brain tissues. Performing a PFA-mediated cross-linking of RNA before sorting, we were also able to minimize RNA leak caused by the sorting procedure. In doing so, we obtained endothelial cell nuclei-specific RNA, as characterized by the enrichment of canonical endothelial cell-specific transcripts. This approach yielded high-quality RNA transcripts from only 100 sorted nuclei, suggesting a potential application to interrogate endothelial cells of the blood brain barrier and other tissues, using bulk or single-cell analysis, to study disease.

## Acknowledgments

We are grateful for the help of Christopher (“Kit”) Bonin and Geneva Hargis in the UCONN Medical Science Writing group for editing.

## Ethics Approval and Consent to Participate

All mice were housed and handled in accordance with IACUC approved protocols in accordance with University of Connecticut Health Center for Comparative Medicine. Human tissue samples were acquired from deceased individuals through the NeuroBioBank maintained by the National Institutes of Health, and passed without patient identifiers to the investigators.

## Author Contributions

P.M. developed the study concept and design; P.M. performed development of methodology. P.M. and J.S. wrote, reviewed and revised the paper; P.M., A.K., J.S. and M.M. provided acquisition of data. P.M. provided analysis and interpretation of data, and statistical analysis; P.M., A.K., J.S., M.M., J.H., S.N., E.J. and B.R. provided technical and material support. All authors read and approved the final paper.

## Data Availability Statement

Key data are provided as a supplementary table. Raw sequencing data will be deposited in dbGaP.

## References

1. Brown, W. R. & Thore, C. R. Review: cerebral microvascular pathology in ageing and neurodegeneration. Neuropathol Appl Neurobiol 37, 56–74 (2011).

2. Habib, N. et al. Massively parallel single-nucleus RNA-seq with DroNc-seq. Nat Methods 14, 955–958 (2017).

3. Mathys, H. et al. Single-cell transcriptomic analysis of Alzheimer’s disease. Nature 570, 332–337 (2019).

4. Darmanis, S. et al. A survey of human brain transcriptome diversity at the single cell level. Proceedings of the National Academy of Sciences of the United States of America 112, 7285–90 (2015).

5. Lake, B. B. et al. Integrative single-cell analysis of transcriptional and epigenetic states in the human adult brain. Nat Biotechnol 36, 70–80 (2018).

6. Ding, J. et al. Systematic comparison of single-cell and single-nucleus RNA-sequencing methods. Nat Biotechnol (2020) doi:10.1038/s41587-020-0465-8.

7. Zhang, Y. et al. An RNA-sequencing transcriptome and splicing database of glia, neurons, and vascular cells of the cerebral cortex. J Neurosci 34, 11929–47 (2014).

8. Munji, R. N. et al. Profiling the mouse brain endothelial transcriptome in health and disease models reveals a core blood-brain barrier dysfunction module. Nat Neurosci 22, 1892–1902 (2019).

9. Vanlandewijck, M. et al. A molecular atlas of cell types and zonation in the brain vasculature. Nature 554, 475–480 (2018).

10. Rosenberg, A. B. et al. Single-cell profiling of the developing mouse brain and spinal cord with split-pool barcoding. Science 360, 176–182 (2018).

11. Bakken, T. E. et al. Single-nucleus and single-cell transcriptomes compared in matched cortical cell types. PLoS One 13, e0209648 (2018).

12. Lacar, B. et al. Nuclear RNA-seq of single neurons reveals molecular signatures of activation. Nat Commun 7, 11022 (2016).

13. Dobin, A. et al. STAR: ultrafast universal RNA-seq aligner. Bioinformatics 29, 15–21 (2013).

14. Sterne-Weiler, T., Weatheritt, R. J., Best, A. J., Ha, K. C. H. & Blencowe, B. J. Efficient and Accurate Quantitative Profiling of Alternative Splicing Patterns of Any Complexity on a Laptop. Mol Cell 72, 187-200.e6 (2018).

15. McKenzie, A. T. et al. Brain Cell Type Specific Gene Expression and Co-expression Network Architectures. Sci Rep 8, 8868 (2018).

16. Zhang, Y. et al. Purification and Characterization of Progenitor and Mature Human Astrocytes Reveals Transcriptional and Functional Differences with Mouse. Neuron 89, 37–53 (2016).

17. Jiang, Y., Matevossian, A., Huang, H. S., Straubhaar, J. & Akbarian, S. Isolation of neuronal chromatin from brain tissue. BMC Neurosci 9, 42 (2008).

18. Krishnaswami, S. R. et al. Using single nuclei for RNA-seq to capture the transcriptome of postmortem neurons. Nat Protoc 11, 499–524 (2016).

19. Birdsey, G. M. et al. The endothelial transcription factor ERG promotes vascular stability and growth through Wnt/beta-catenin signaling. Developmental cell 32, 82–96 (2015).

20. Haber, M. A. et al. ERG is a novel and reliable marker for endothelial cells in central nervous system tumors. Clin Neuropathol 34, 117–27 (2015).

21. Martowicz, A. et al. Endothelial beta-Catenin Signaling Supports Postnatal Brain and Retinal Angiogenesis by Promoting Sprouting, Tip Cell Formation, and VEGFR (Vascular Endothelial Growth Factor Receptor) 2 Expression. Arteriosclerosis, thrombosis, and vascular biology 39, 2273–2288 (2019).

22. Nagai, N. et al. Downregulation of ERG and FLI1 expression in endothelial cells triggers endothelial-to-mesenchymal transition. PLoS Genet 14, e1007826 (2018).

23. Murphy, P. A. et al. Alternative RNA splicing in the endothelium mediated in part by Rbfox2 regulates the arterial response to low flow. Elife 7, (2018).

24. Boutz, P. L., Bhutkar, A. & Sharp, P. A. Detained introns are a novel, widespread class of post-transcriptionally spliced introns. Genes Dev 29, 63–80 (2015).

25. La Manno, G. et al. RNA velocity of single cells. Nature 560, 494–498 (2018).

26. Evers, D. L., Fowler, C. B., Cunningham, B. R., Mason, J. T. & O’Leary, T. J. The effect of formaldehyde fixation on RNA: optimization of formaldehyde adduct removal. J Mol Diagn 13, 282–8 (2011).

27. Attar, M. et al. A practical solution for preserving single cells for RNA sequencing. Sci Rep 8, 2151 (2018).

28. Liu, E. Y. et al. Loss of Nuclear TDP-43 Is Associated with Decondensation of LINE Retrotransposons. Cell reports 27, 1409–1421 e6 (2019).

29. Stoeckius, M. et al. Simultaneous epitope and transcriptome measurement in single cells. Nat Methods 14, 865–868 (2017).

