## Supplemental Table 1 for "A Method for Rapid Flow-cytometric Isolation of Endothelial Nuclei and RNA from Archived Frozen Brain Tissue"

| *Group* | *Sex* | *Age* | *PMI* |
|---|---|---|---|
| Young | M | 19 | 13 |
| Young | M | 20 | 18 |
| Young | M | 22 | 13 |
| Old | M | 68 | 28 |
| Old | M | 69 | 23 |
| Old | M | 71 | 18 |
| Dementia | M | 58 | 3 |
| Dementia | M | 72 | 5 |
| Dementia | M | 74 | 14 |
| Dementia | M | 65 | 12 |
| Dementia | M | 67 | 16 |
| Old-test | F | 87 | 4 |

**SI Table 1. Archived human brain tissue from NIH NeuroBioBank**
Frozen cortical tissues (Brodmann Area 10) were obtained from each of the indicated samples. Tissues were cut into 200mg pieces, and separately analyzed by staining and flow-cytometry using a protocol with and without brief 4% PFA fixation.
