## Supplemental Figures for "A Method for Rapid Flow-cytometric Isolation of Endothelial Nuclei and RNA from Archived Frozen Brain Tissue"

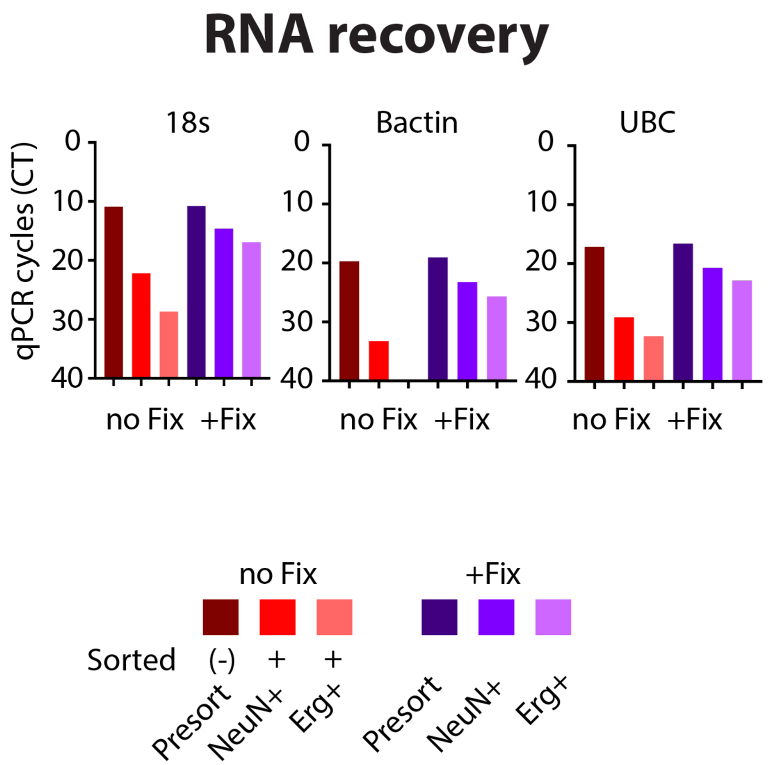


**Supplemental Figure 1. Fixation of RNA in nuclei allows improved recovery after sorting.** Recovery of RNA from unsorted nuclei from mouse brain (200mg tissue), and from nuclei after sorting, with or without fixation. 10% of the nuclei from the tissues were used to prepare cDNA for the unsorted fraction and all of the nuclei were used to prepare cDNA for the sorted fractions. Graphs show the threshold cycle (CT) from quantitative PCR analysis of the indicated genes.

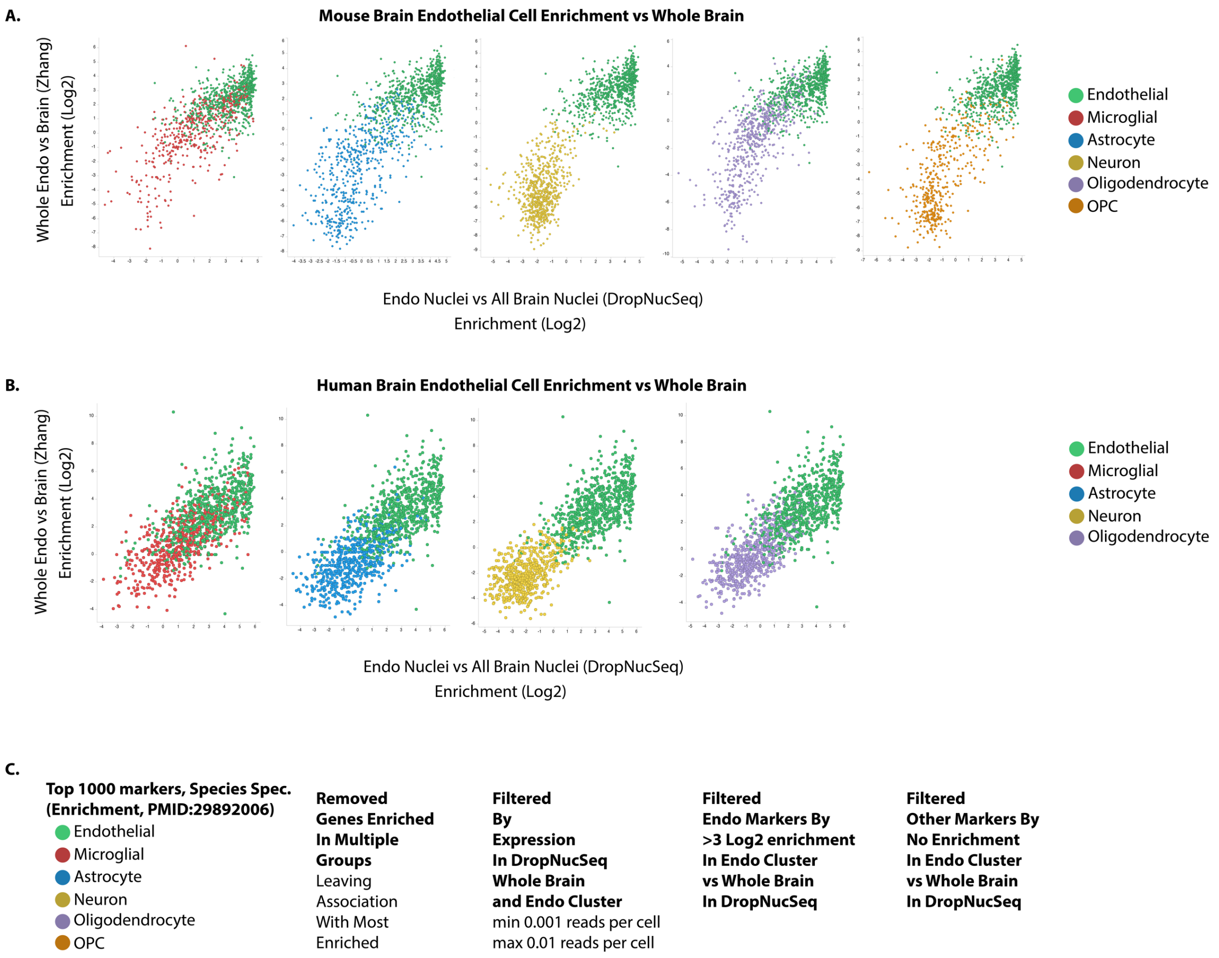


**Supplemental Figure 2. Cell markers used in enrichment analysis.**
Plots show the endothelial enrichment of individual genes (points on the plot), for genes previously identified as enriched in the indicated cell types from a meta-analysis of cell-specific RNA sequencing experiments in (A) mouse and (B) human brain tissues. Datasets used to plot enrichment were (A) Zhang whole endothelial cell isolation vs. whole mouse cortex (PMID: 25186741) and DropNucSeq data from endothelial cluster 21 vs. all nuclei (all clusters) (PMID: 28846088) and (B) Zhang whole endothelial cell isolation vs. bulk human cortex (PMID: 26687838) and DropNucSeq data from endothelial cluster 16 vs. all nuclei (all clusters) (PMID: 28846088). (C) The marker panel was trimmed as described for mouse and human markers individually, resulting in the pared list of genes used as markers for enrichment analysis.

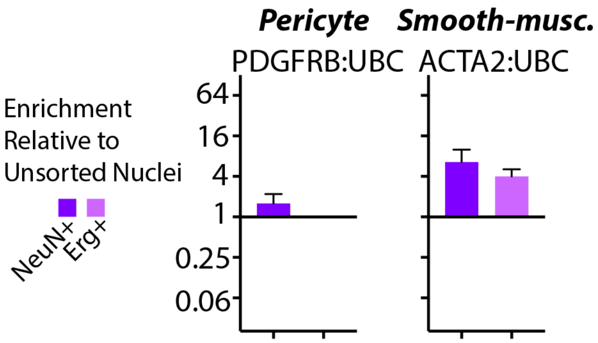


**Supplemental Figure 3. Mural cell markers are not enriched in the Erg+ fraction.** Enrichment of cell type-specific RNA from the sorted Erg+ or NeuN+ nuclei versus unsorted nuclei by quantitative PCR. The fold-change increase in each transcript, relative to unsorted nuclei is shown.

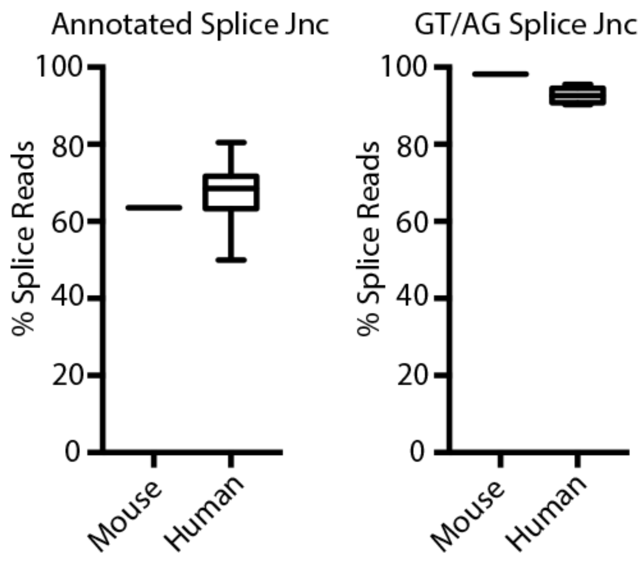


**Supplemental Figure 4. Annotation of splice junction reads from Erg+ samples.**
Graphs show the STAR analysis of splice junction reads, and the percentage mapping to known splice junctions in the Hg38 annotation file, and those with GT/AG junction (canonical splice junction dinucleotides).
